# RNAcentral in 2026: Genes and literature integration

**DOI:** 10.1101/2025.09.19.677274

**Authors:** Andrew Green, Carlos Eduardo Ribas, Isaac Jandalala, Philippa Muston, Colman O’Cathail, Guy Cochrane, Christina Ernst, Lingyun Zhao, Pedro Madrigal, Helen Attrill, Steven Marygold, Doron Lancet, Niv Dobzinski, Patricia P. Chan, Todd M. Lowe, Elspeth A. Bruford, Ruth L. Seal, Henning Hermjakob, Kalpana Panneerselvam, Robert D. Finn, Tatiana A. Gurbich, Sam Griffiths-Jones, Bastian Fromm, Kevin J. Peterson, Dominik Sordyl, Janusz M. Bujnicki, Sameer Velankar, Sri Devan Appasamy, Sudakshina Ganguly, Peng Zhang, Shunmin He, Kim M. Rutherford, Valerie Wood, Ruth C. Lovering, Ernesto Picardi, Nancy Ontiveros, Lin Huang, Zhichao Miao, Anton S. Petrov, Holly McCann, Emanuele Cavalleri, Marco Mesiti, Elena Rivas, Marcell Szikszai, Marcin Magnus, Jan Gerken, Maria Chuvochina, Danny Bergeron, Michelle Scott, Kelly Williams, Robin R Gutell, Cheong Xin Chan, Mark Quinton-Tulloch, Stavros Diamantakis, Anton I. Petrov, Alex Bateman, Blake A. Sweeney, The RNAcentral Consortium

## Abstract

RNAcentral was founded in 2014 to serve as a comprehensive database of non-coding RNA sequences. It began by providing a single unified interface to more specialised resources, and now contains 45 million sequences. It has grown beyond providing a single interface to many specialised resources and now provides several services and analyses. These include secondary structure prediction with R2DT, sequence search, and analysis with Rfam. Since its last publication in 2021, RNAcentral has developed two major features. First, literature integration with the development of LitScan and LitSumm. LitScan automatically identifies and links relevant publications to RNA entries, while LitSumm uses natural language processing to generate functional summaries from the literature. Together, these tools address the critical challenge of connecting sequence data with scattered functional knowledge across thousands of publications. Secondly, RNAcentral has created gene level entries. Gene level entries represent a large structural change to RNAcentral. While RNAcentral previously organized data exclusively at the sequence level, we now group related transcripts into gene-centric views. This allows researchers to explore all isoforms, splice variants, and related sequences for a gene in a unified interface, better reflecting biological organization and facilitating comparative analyses. RNAcentral is freely available at: https://rnacentral.org.

## Introduction

RNAcentral was established in 2014 as a comprehensive database unifying all non-coding RNA (ncRNA) sequences into a single, searchable resource (1). This was aimed at solving the issue of fragmentation in the non-coding RNA field, where there were many high quality databases that focused on specific subtypes of ncRNA. There was no easy to use resource that compiled all ncRNA data and RNAcentral sought to fill that gap. RNAcentral is designed around an ‘expert database’ model, where each contributing database provides its expertise and data in a single area, and then RNAcentral combines, standardizes and integrates these data into a single resource. Since its inception it has grown to contain over 45 million sequences and has moved from providing sequences and metadata to more complex data types, such as Gene Ontology (GO) annotations (2), Sequence Ontology (SO) terms (3), interaction data from resources such as IntAct (4) and RNA-KG (5, 6), and analysis, such as Rfam and R2DT predictions (7). RNAcentral remains the primary source of sequence data for ncRNA science, with tens of thousands of users and hundreds of citations per year.

Since its last publication in 2021 (7), RNAcentral has had 9 releases, grown from 30 to 45 million sequences and now includes 52 expert databases, including ten new databases and major updates to existing sources. We also expanded secondary structure predictions from 13 to 30 million sequences. Beyond these quantitative changes we have also updated and improved how researchers can interact with our ncRNA data. We extended our search to allow searching by taxonomic descendants (e.g., all primates or all fungi), and transitioned to CC0 licensing for unrestricted data reuse enabling both academic and commercial reuse.

This article details the advances in RNAcentral release 26: new database integrations and major updates to existing resources; the implementation of our literature integration system (LitScan and LitSumm); the creation of gene level entries; technical infrastructure improvements; and future development priorities.

## Database Content and Growth

Since our 2021 publication, RNAcentral has grown from 30 to 45 million sequences (50% increase) across 52 expert databases. This expansion includes 10 new database integrations, major updates to three core resources, and the retirement of one database, VEGA. Table 1 summarizes the major changes across releases 17-26.

**Table 1:** A summary of the key features added since the last publication (releases 17-26).

| Release | Sequences (millions) | Key features |
| --- | --- | --- |
| 17 | 30 | Added piRBase, added Ribovore |
| 18 | 30 | Finalized improved rRNA RNA types |
| 19 | 31 | Added PSICQUIC, CPAT based ORF prediction, |
| 20 | 31 | Migrated to CC0, Added RiboVision, Created LitScan |
| 21 | 33 | Added PLncDB, Added Expression Atlas, Added SwissBioPics |
| 22 | 34 | Added EVlncRNAs, Added Ribocentre, Integrated Expression Atlas Viewer |
| 23 | 36 | Added MGnify, Added REDlportal, Improved genome browser, Added LitSumm |
| 24 | 35 | Added tmRNA Website 2.0, Improved Phylogenetic search |
| 25 | 45 | Added Rfam 15.0, Improved taxonomic search |
| 26 | 45 | Created gene level entries, Added RNA-KG |

### New Database Integrations

We integrated ten new expert databases to expand both taxonomic coverage and functional annotations. piRBase (8) contributes 200,000 piRNA sequences from 21 organisms for studying transposon silencing. PSICQUIC (9) provides 4,420 manually curated RNA-protein interactions to expand our coverage of RNA interactions. PLncDB (10) adds plant lncRNAs from 80 species, while linking to EVlncRNAs (11) provides a connection to their experimentally-validated lncRNAs from 19,000 publications with disease associations. Ribocentre and Ribocentre-switch (12, 13) provide structural data for ribozymes and riboswitches respectively, enabling structure-function studies. MGnify (14) expands environmental coverage with sequences from over 28,000 metagenome-assembled genomes spanning 1,132 isolates. RiboVision (15) provides templates to R2DT, allows secondary structure editing and connects sequences to 3D structures and detailed annotations. REDIportal (16) adds RNA editing events for post-transcriptional modification analysis. Expression Atlas (17) provides tissue-specific expression profiles with interactive visualizations, expanding sequence data with functional information. RNA-KG (5, 6) integrates around 100M interactions involving RNAs from 91 linked open data repositories and ontologies. Relationships are characterized by standardized properties that capture the specific context (e.g., cell line, tissue, pathological state) in which they have been identified.

A key database RNAcentral now incorporates is Expression Atlas (17). This resource hosts gene and protein expression at bulk and single cell levels for a range of species. The data provided is often at the tissue level. RNAcentral has imported the bulk expression data and mapped their gene level entries to our sequence pages. We have also integrated the visualisation provided by Expression Atlas to allow users to explore the expression of their sequence of interest.

Overall, RNAcentral increased the range of ncRNAs included, adding piRNAs with piRBase, expanded to new organisms, e.g. plant and metagenome from PLncDB and MGnify, but also enriching the data included by adding expression and modification data via Expression Atlas and REDIportal. Users interested in exploring data from any of these resources can use our text search to filter by database.

### Major Database Updates

In addition to adding new resources, we have also had several resources undergo major changes. First, Rfam updated to version 15.0 (18). This was a major update to Rfam which included updating the matches to all families. This led to increasing the size of Rfam from 1.88M to 5.9M sequences, with refreshed genomes and updated alignments for all RNA families, substantially improving family coverage and annotation quality. Secondly, TarBase is now at version 9.0 (19). TarBase now contains experimentally validated miRNA-gene interactions from a range of species, including virally encoded miRNAs. Finally, we are now the primary site hosting Transfer-messenger (tmRNA) Website data (20). The site has been discontinued, but the data has been updated in RNAcentral. This now expands RNAcentral to include an archival role for expert databases, where the data can continue to be publicly accessible after resources become unavailable. In release 24, we updated the data and included detailed annotations of sequence features within tmRNAs (21).

### Enhanced Analysis Pipelines

Since release 17 RNAcentral has added two new analyses. First, we now use Ribovore to analyze all sequences from ENA (1) prior to import (22). Ribovore is a tool which can quickly and accurately detect complete and fragmentary rRNAs. We found that the rate of fragmentary rRNAs was doubling the size of RNAcentral but provided little value to users as these are generally poorly annotated sequences. Thus we sought to exclude them from future import and used Ribovore to detect such sequences. This does not prevent the import of complete rRNAs, or non metagenomic rRNAs.

In addition, we now analyse sequences from 4 organisms, human, fly, zebrafish and mouse, with CPAT, which detects open reading frames (23). Results indicate that approximately 6% of sequences contain detectable ORFs, providing quality metrics that help distinguish coding transcripts from legitimate non-coding RNAs. These sequences are given a QC warning as containing an open reading frame. Finally, we used the existing Rfam analysis pipeline to reanalyze all sequences with all Rfam families after each major release of Rfam. This will keep the annotations up-to-date with the latest families and metadata.

## Literature Integration: Connecting RNA Sequences with Scientific Knowledge

A longstanding challenge for RNAcentral has been providing users with comprehensive, up-to-date connections between RNA sequences and their functional descriptions in the scientific literature. To address this need, RNAcentral has developed two complementary tools to automatically extract and synthesize information from the vast corpus of RNA research: LitScan for literature mining and LitSumm for automated summarization (24) that work with any ncRNA in RNAcentral.

### LitScan: Mining the Literature for RNA Mentions

LitScan systematically identifies and extracts literature mentions of RNA identifiers in the open access literature. The system queries the EuropePMC (2) API using RNAcentral’s comprehensive collection of identifiers and their aliases.

For each matching article, LitScan extracts the complete sentences with the mentioned RNA. It uses RNAcentral’s cross-references to capture alternative names, for example the lncRNA THRIL is also known as “Linc1992” and “NR_110375”. Results are stored in RNAcentral’s database and displayed through an interactive widget allowing filtering by journal, publication year, article section, and publication type (Figure 1). Additionally, some expert databases provide annotations for key papers that discuss an RNA. RNAcentral captures this information and provides it as a filter in the viewer.

**Figure 1.**
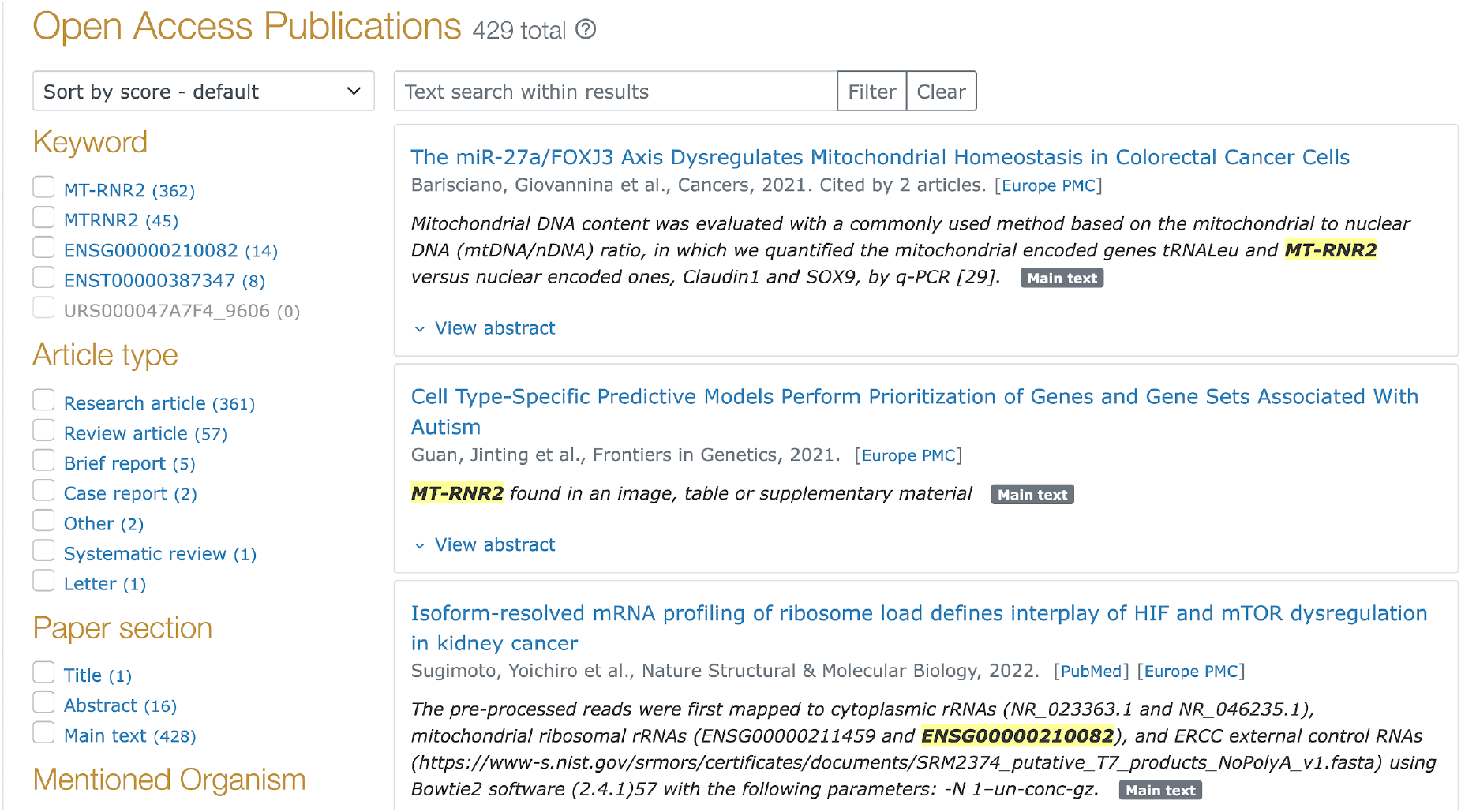
Image of an RNAcentral sequence page for human MT-RNR2 (URS000047A7F4_9606) with the LitScan widget displayed.

As of release 26, RNAcentral has processed 40 million papers with 13.6 million identifiers, resulting in 1.5 million ncRNA sequences connected to at least one paper. Most RNAs have very few papers with well-studied RNAs such as human XIST and MALAT1 appearing in over 5,000 publications.

### LitSumm: LLM-Powered Literature Summarization

LitSumm addresses the challenge of synthesizing scattered functional knowledge by generating structured summaries from LitScan-extracted sentences. The system employs a multi-stage pipeline, where sentences from LitScan are selected via topic modeling, an initial summary is generated using ChatGPT4 and then checked for self-consistency and sufficient citations (24). We then utilize GPT-4-turbo (gpt-4-1106-preview) to summarise and validate the summaries. Expert review of summaries showed that 94% of the summaries were rated good or excellent. Where the summary was rated poorly, the main reason was a failure to properly synthesize facts from multiple sources. This issue, which automated checking could not catch, has also been observed in other work (25). The other most common issue was references being either irrelevant or assigned to the incorrect sentence. A detailed analysis of the limitations of LitSumm is given in (24).

Currently, LitSumm has generated summaries for approximately 4,600 human ncRNAs prioritized by community interest (HGNC (3), miRBase (4), mirGeneDB (5), and snoDB (6) entries). For miRNAs, we restrict to species-specific identifiers (e.g., “hsa-mir-21” rather than “mir-21”) to prevent cross-species confusion. The LitSumm summaries are displayed on each sequence page accompanied by a warning about AI generated content and a link to provide feedback (Figure 2). Entries with summaries are available on RNAcentral by searching for ‘has_litsumm:”True”‘(https://rnacentral.org/search?q=has_litsumm:%22True%22) or can be fetched at: https://huggingface.co/datasets/RNAcentral/litsumm-v1.5 and the code is available at: https://github.com/RNAcentral/litscan-summarization.

**Figure 2.**
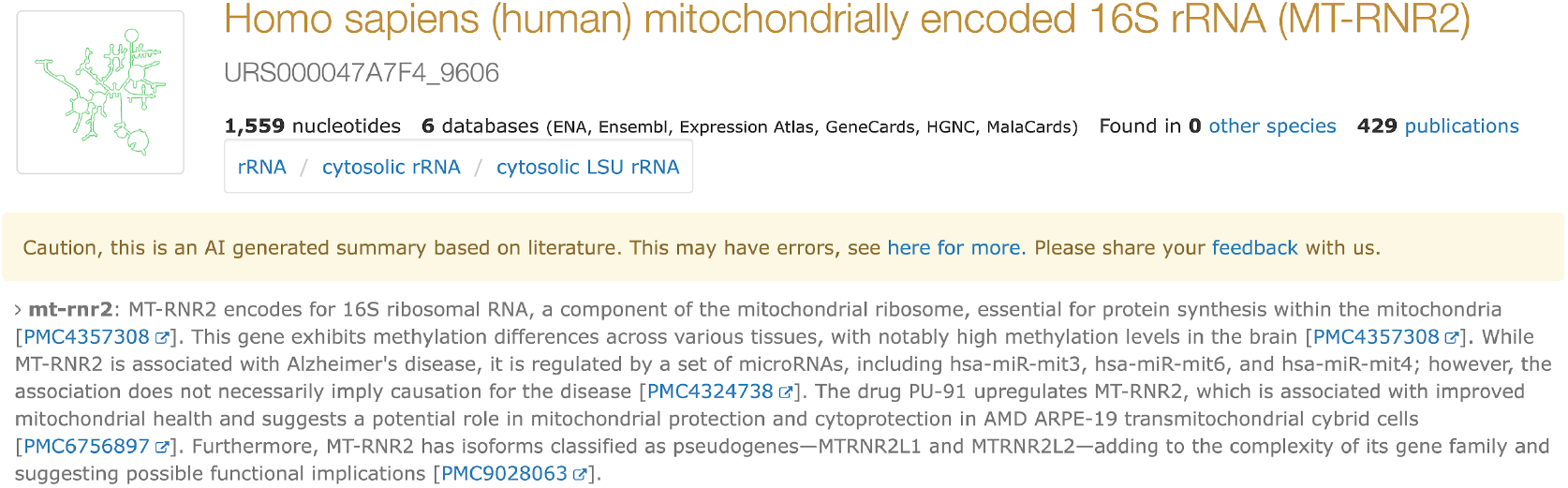
Image of an RNAcentral sequence page for human MT-RNR2 (URS000047A7F4_9606), with the LitSumm summary highlighted.

**Figure 3.**
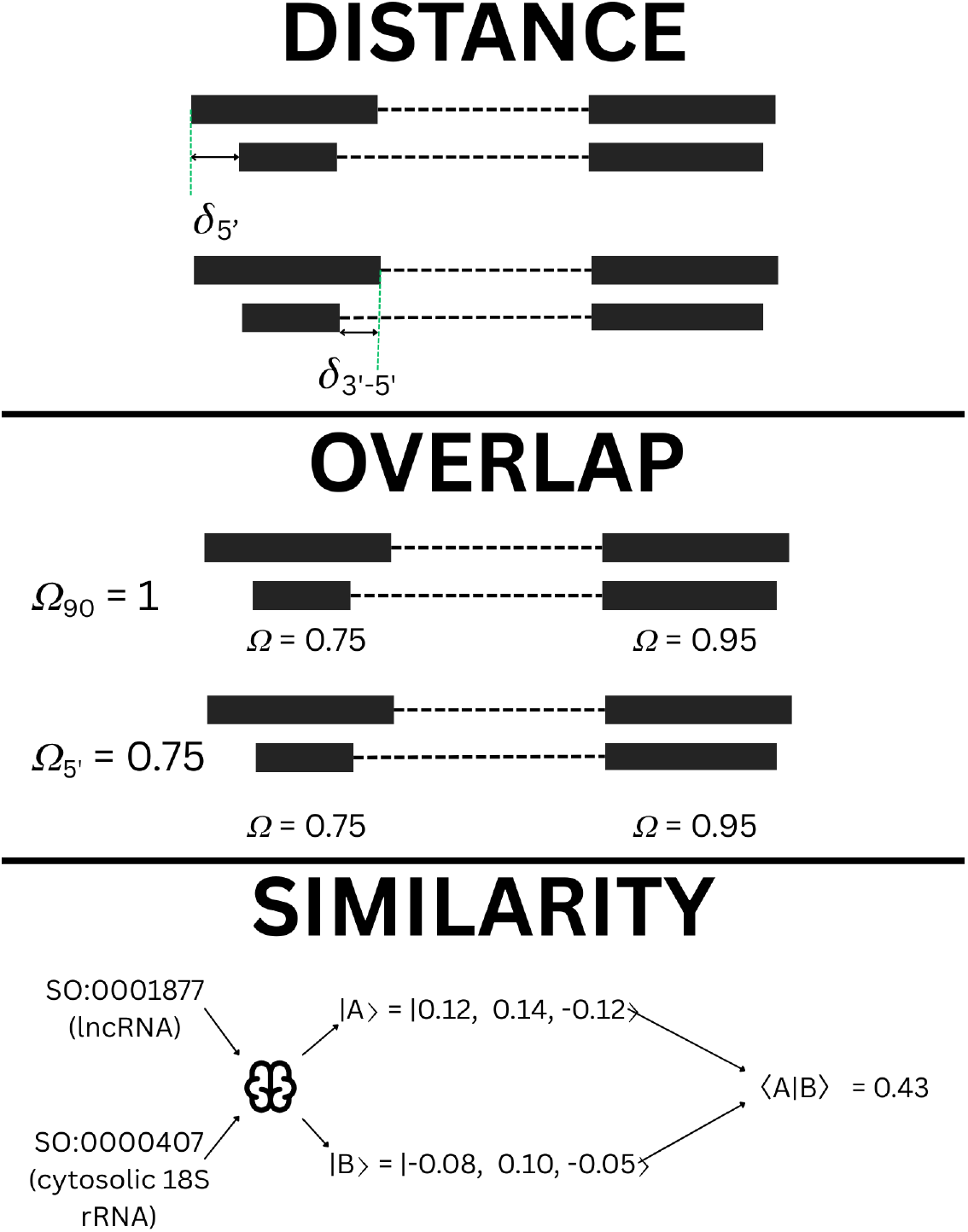
Illustration of feature calculation based on mapped transcript coordinates. Distance metrics primarily focus on the 5’ exon, and are simply the distance in number of nucleotides between the two transcripts. Overlap metrics consider the 5’ exon as well as the global exon structure, measuring the overlap of the 5’ exons of two transcripts, and the number of exons within the transcripts having >90% overlap. Finally, the type similarity uses a Node2Vec model trained on the sequence ontology terms to convert types into normalised vectors, then calculates the inner product to give a numerical similarity between RNA types.

Development of LitSumm is ongoing, aiming to alleviate the limitations of the original. Among these is the sensitivity of the original system to mentions ‘in passing’ where a paper merely mentions an ncRNA but is not the main focus of the study. We are developing an agentic approach including literature triaging and prioritisation to filter out such mentions, and provide much better initial context for the summary. The second version of LitSumm will continue to prioritise information provenance above all else, and is expected to enter production in mid 2026.

### RNAcentral gene building

In release 26 of RNAcentral we have created the first set of genes for all sequences which could be found in a genome, excluding piRNAs, in 204 organisms. Until now RNAcentral has been a sequence-based resource, which means each unique sequence is given a Unique RNA Sequence identifier (URS id) (1) and treated as separate entries. There are many examples of sequences which may differ in only a few nucleotides. When these sequences are present in the genome, they often represent variants in sequence of the same transcript. For example, they include cases of one transcript a single nucleotide longer than another being treated as separate unrelated entries. For genes that are commonly used for fingerprinting, such as rRNAs, there will be thousands of examples of sequences which differ slightly in overall length and have a few changes between them. Conversely, for some ncRNAs, such as miRNAs, there may be several identical copies in a genome, which are merged into a single URS entry. Both of these situations can also be confusing to users. Additionally, many biological experiments and data are at the gene, not transcript level. These issues have led us to build gene-level entries into RNAcentral.

We could not simply use gene objects from existing resources, such as Ensembl (7), because RNAcentral’s data includes many sequences which are absent from other collections. However, the goal was to produce gene objects which are comparable to those in Ensembl for all organisms in RNAcentral without human intervention. Finally, the gene building pipeline must be able to accommodate the changes that occur between RNAcentral releases.

As each RNAcentral release may add thousands of new sequences to an organism the pipeline must be able to detect when two genes in two different releases are similar enough to be revised versions of the same gene. RNAcentral genes should have stable identifiers across releases even in the presence of changes. We note that in RNAcentral a single release adds around 1 million new sequences and may add several thousand new ones to highly studied organisms such as humans. The pipeline must tolerate large changes in the number of transcripts within a gene.

### Gene building pipeline

The new RNAcentral gene building pipeline solves these problems using two major steps. First, for each release we run a graph clustering algorithm to build transcript clusters, and then we compare transcript clusters to the previous release to assign gene identifiers. Each transcript is treated as a node, and all pairs of transcripts, whose start site is within 1kb, are compared using a random forest model to determine which pairs should be connected. We then find communities (26) in the resulting graph and treat those communities as genes. Communities were used to prevent the tendency of rare but large transcripts to join several otherwise distinct genes. We exclude piRNAs from this gene building effort, as these ncRNAs have a unique biogenesis pathway (27).

To guide the development of the random forest model we asked approximately 30 consortium members to manually cluster 10-20 transcripts in 3 example regions. We then discussed with them why certain transcripts were clustered together and what features they considered. We used these features and trained a random forest on the existing Ensembl/GENCODE (8) transcripts from the human genome. We selected the human genome as this has undergone the most manual curation. We use three classes of measures, distance, overlap, and similarity. The distance measures the proximity of the start and end of the 5’ most exon. For overlap we look at how many exons have more than 90% overlap, and the percentage overlap of the 5’ most exon. Finally for similarity we use node2vec (28) to compute similarity between the SO RNA types between transcripts as assigned by the source expert databases. Node2vec is a method which allows the calculation of vector similarity within a graph structure. As the SO is a directed acyclic graph this allows us to easily compute similarity between any pair of SO terms.

The random forest model is used to infer the probability that any given pair of transcripts come from the same gene. This pairwise comparison is then used to provide an edge weight between nodes in the graph, which is used when calculating the communities that become genes.

We used 5 fold cross validation when training the procedure on human genes and found excellent performance, with an average Area Under the Receiver-Operator Curve (AUC) 0.994 (Table 3) - where an AUC of 1.0 would be a perfect classifier. As this performance could be due to imperfect splits in the data, we applied the model to Ensembl annotations of a similar well annotated species, mouse, and observed consistently high performance (Average AUC 0.974, Table 4).

**Table 3:** Cross validation performance of the gene building model built and tested on human genes from Ensembl.

| **Fold** | **Accuracy** | **F1** | **AUC** | **AP** |
| --- | --- | --- | --- | --- |
| 1 | 0.968 | 0.989 | 0.994 | 0.998 |
| 2 | 0.970 | 0.989 | 0.995 | 0.998 |
| 3 | 0.970 | 0.989 | 0.995 | 0.998 |
| 4 | 0.968 | 0.989 | 0.994 | 0.998 |
| 5 | 0.968 | 0.989 | 0.994 | 0.997 |
| **Average** | 0.969 | 0.989 | 0.994 | 0.997 |

**Table 4:** Cross validation performance of the gene building model built on human genes but on mouse genes from Ensembl.

| **Fold** | **Accuracy** | **F1** | **AUC** | **AP** |
| --- | --- | --- | --- | --- |
| 1 | 0.887 | 0.959 | 0.975 | 0.989 |
| 2 | 0.887 | 0.959 | 0.975 | 0.989 |
| 3 | 0.887 | 0.959 | 0.975 | 0.990 |
| 4 | 0.888 | 0.960 | 0.976 | 0.990 |
| 5 | 0.888 | 0.960 | 0.969 | 0.980 |
| **Average** | 0.887 | 0.959 | 0.974 | 0.988 |

Inspecting the random forest model, we find the feature importances behave as expected from our discussions with consortium members (Figure 4); the most important features are to do with the distances between 5’ coordinates of the transcripts, both the simple distance between the two 5’ transcript start coordinates, but also the distance between the coordinates of the 3’ end of the 5’ exon. Transcripts being located on the same strand was obviously also important, but sequence ontology type similarity was less crucial than we expected. The relative unimportance of sequence type similarity may be a reflection of the training data used (coming from known genes) having relatively low diversity.

**Figure 4.**
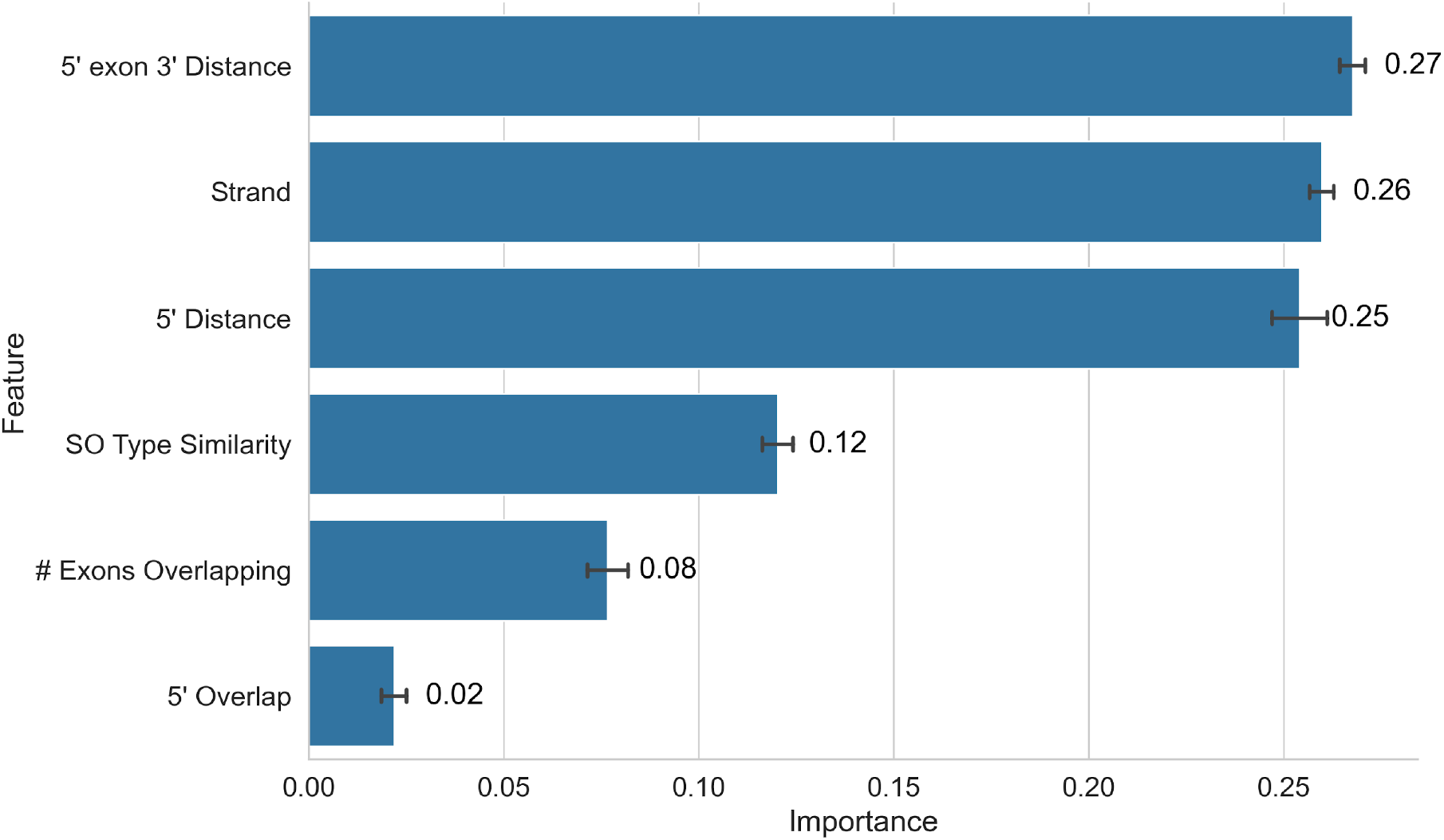
Feature importances from the random forest model, averaged across the five folds of cross validation. Feature importances were calculated using the mean decrease in gini impurity, aggregated over all trees in the forest, with confidence intervals derived from the standard deviation of importance across folds. These match what was expected from discussion with experts, with measures of distance at the 5’ end of transcripts being most informative, alongside the transcripts being mapped to the same strand. SO type similarity was relatively unimportant for the model, possibly reflecting a lack of diversity in the types contained in the training data.

We then developed logic to merge clusters between releases. Because the build pipeline uses only data available from RNAcentral’s GFF files we were able to run it for all sequences from release 12, from September 2019, onward. This allowed us to inspect how different merging criteria would change the resulting genes. After inspecting the data, we found that treating clusters with a start site within 1kb between two releases as being the same gene produced stable results.

### Gene identifiers and metadata

RNAcentral genes are given accessions with the following pattern ‘RNACG<species-prefix><11-digit hash>.<version>’. This allows tracking the organism, identity, and version of these genes over time. The species prefix is the same as the Ensembl species prefix, and for *Homo sapiens* there is no species prefix. The 11 digit hash is computed using sha256 from the gene coordinates, chromosome and assembly ID and is intended to be similar to gene names from Ensembl. The version number of each gene increments by one each time there is any change to the gene. These changes include adding or removing transcripts, or changing the start or end points of a gene. If no changes are made, then a gene will retain the same accession between releases. Some genes have remained unchanged since creation in release 12, while others have changed with every release.

Alongside the identifier, each gene is given an RNA type based on SO terms and a description. These are assigned from the RNA types and descriptions provided by expert databases and the Rfam and R2DT annotations. The R2DT template assignments supplemented Rfam family annotations to identify specific SO types, such as tRNA and rRNA types. We used a weighted voting system where data from curated resources such as HGNC (3), FlyBase (9) and GENCODE are weighted more than non-curated resources such as ENA. Similarly, we only consider results from Rfam or R2DT if they match over 90% of the longest sequence in the gene. Manual inspection of the gene names and types indicates this system performs well overall, however, we plan to continue to improve and extend the system to better deal with edge cases and uncertainty.

### Current human gene set

RNAcentral contains 103,814 human non-coding RNA genes from 600,225 transcripts. These genes cover 56 different SO terms, with the most common being lncRNA, 65,187, antisense lncRNA, 16,790, followed by pre-miRNA, 8,560. The remaining 13,277 genes are SRP RNA, generic ncRNAs and a variety of specific terms, e.g. tRNA subtypes. The average ncRNA gene in RNAcentral contains 6 transcripts, with the minimum being 2 and maximum being 4,001 for a mitochondrial small subunit rRNA with many fragments. Comparing the genes to Ensembl shows that or on average RNAcentral genes map to 1.6 Ensembl genes, indicating our pipeline, will on average merge genes that Ensembl considers separate.

We examined several specific cases, including MALAT1, NEAT1, XIST the clustering of a miRNA sequence URS000075CC93_9606. We found that both MALAT1 and NEAT1 were each built into a single gene as expected. On the other hand, the miRNA URS000075CC93_9606 represents several different miRNAs, hsa-mir-1302-2 and hsa-mir-1302 9 to 11. Ensembl and miRBase list this sequence as occurring in 4 different genes, and RNAcentral’s gene building reflects that. On the other hand, some genes are not built correctly. For example XIST, ENSG00000229807, is split across 5 different genes in RNAcentral. This is due to XIST’s complex splicing structure where there is a large distance between possible 5’ exons of XIST. In general, the RNAs which are not built correctly are lncRNAs with complex this alternative splicing pattern. We are looking into correcting this issue, as well as alternative approaches to solving this problem.

RNAcentral provides access to the genes in several ways. First the text search now includes genes, users can select the ‘Genes’ value in the Entry Type facet and see only gene level results. Users are able to use any facet Additionally, all sequence pages link to the genes, if any, the sequence is a part of. Finally, genes have been added to our GFF files as a ‘predicted_gene’ entry.

### Continued Development

The gene pipeline performs well overall, however there is scope for improvement. The identified feature importances indicate that the model will group ncRNA clusters into single genes, rather than as might be expected grouping each ncRNA into its own gene. ncRNA clusters occur for several types including miRNA, tRNA and snoRNA; we are working with type-specific expert databases to refine the gene model for these types. Another limitation arises for the ‘inside-out’ genes such as SNHG1, where the ncRNA product is coded in an intron. For these RNAs, our model is unlikely to perform well, due to limitations in the training data. For training, we use genes and their transcripts from Ensembl, but these do not split ncRNAs in transcript introns into their own transcript; as a result the random forest model has not seen how to handle these correctly. We are working to improve the preprocessing of our training data to remove this limitation.

The set of genes identified by the system now can be considered a superset of the ncRNA genes that exist; this method has high recall, but relatively low precision. While we cannot know for sure which of the predicted genes are real and therefore cannot train directly to improve precision, we can apply further methods to improve precision by filtering out predictions that are demonstrably wrong. We are exploring the use of sequence homology within the predicted gene set to filter out genes we predict that have no evidence of evolutionary conservation; we expect this additional filtering will remove many genes, and improve the quality of those that remain. Alongside these post-prediction filtering steps, we are implementing measures to exclude transcripts with excess protein coding potential from being considered for gene building by incorporating additional QA steps at import, such as Pfam, stopFree (10) and tcode (10).

## Technical Infrastructure and Tools

Since its last publication, RNAcentral has improved several key components: we extended phylogenetic search functionality, migrated to a modern genome browser, updated R2DT to version 2.1, integrated SwissBioPics visualization, and incorporated the Expression Atlas viewer.

### Search and Discovery Enhancements

In release 24, RNAcentral extended phylogenetic search functionality to include subspecies filtering. Users can now toggle between searching only the selected species (e.g., *Escherichia coli*, NCBI taxonomy ID 562) or including all subspecies variants. Now, the user is able to select ‘Include subspecies’ to also find subspecies of *E. coli*. An example of using this feature is shown in Figure 5. This enhancement addresses user requests for more granular taxonomic control and has been applied to all taxonomic entries in RNAcentral.

**Figure 5.**
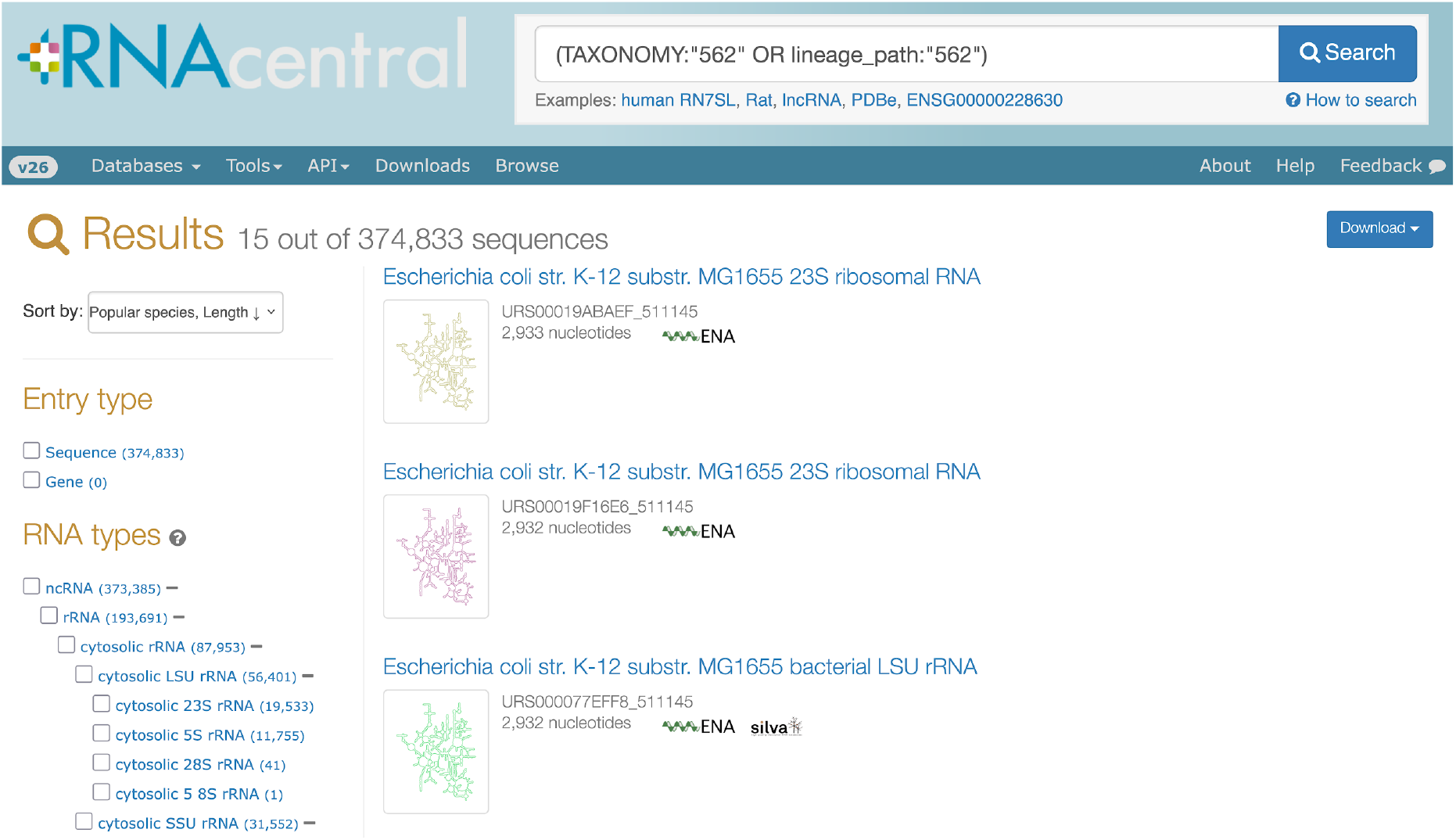
An example of search for all subspecies of *Escherichia coli* in RNAcentral. https://rnacentral.org/search?q=(TAXONOMY:%22562%22%20OR%20lineage_path:%22562%22)

### Improved visualisation

We have improved several aspects of our visualisations. First, our genome browser was migrated from Genoverse to igv.js (Figure 6B) (29). This modern genome browser allows user track uploads, is faster, less error prone and easier to develop with.

**Figure 6.**
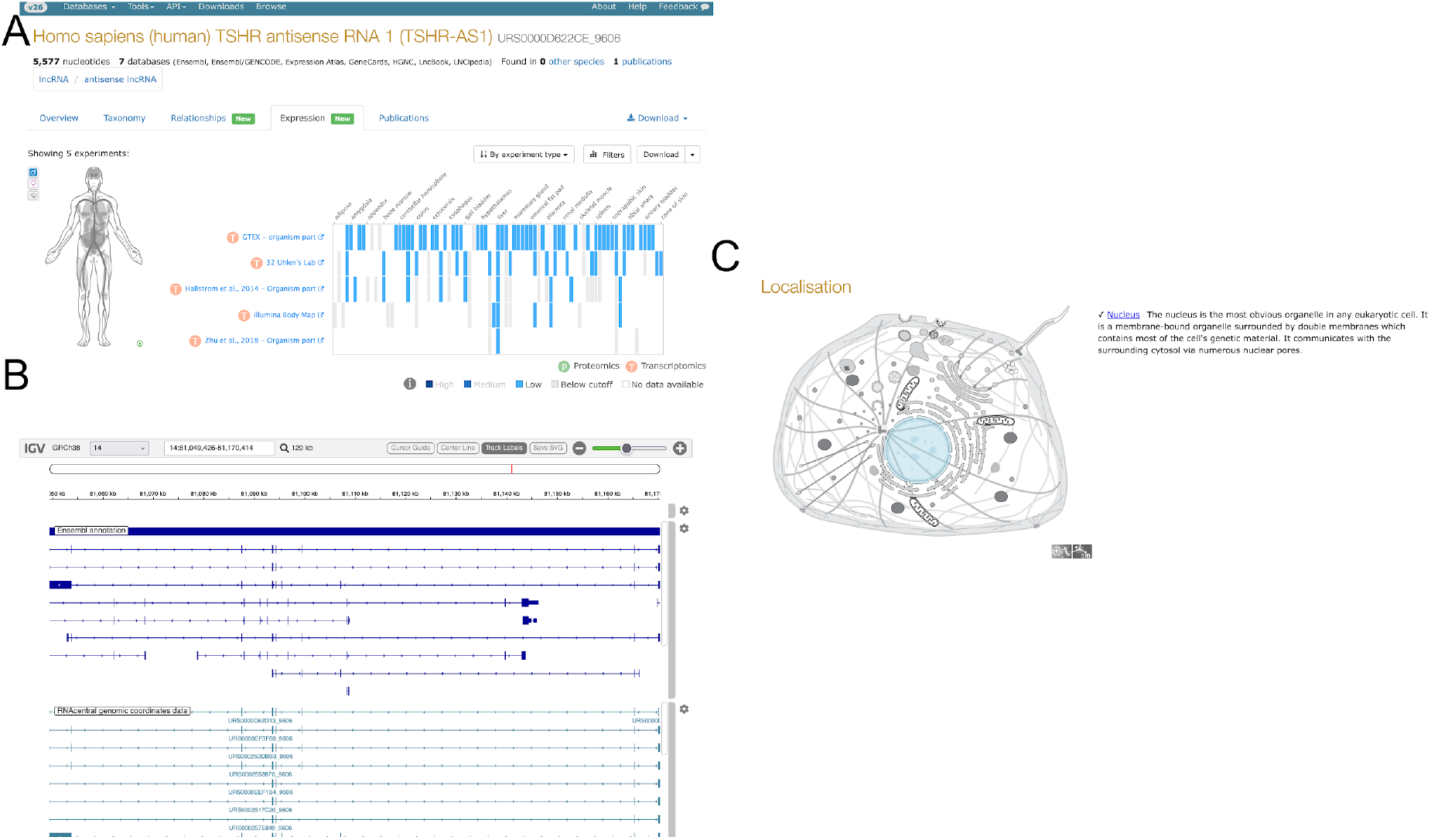
Examples of visualization improvements in RNAcentral. A) The expression atlas viewer for TSHR-AS1 (URS0000D622CE_9606). B) The igv.js based genome browser for TSHR-AS1 (URS0000D622CE_9606). C) The annotated localization of SNORA73A (URS00000DE3E2_9606).

R2DT version 2.1 includes several new templates including mitochondrial tRNAs, tmRNAs and new rRNA models (30). As a result of the update our coverage increased to 30 million sequences from 13 million previously, now covering 55% of sequences.

Third, we have added SwissBioPic integration, which is a visualisation of the subcellular localisation of molecules (Figure 6C) (31). The locations use the ‘located_in’ Gene Ontology annotations imported into RNAcentral. This provides users a simple way to see where a molecule is found in the cell.

The Expression Atlas integration queries data for RNA expression across 4,000+ studies and 60+ species as represented in Expression Atlas. Users can access tissue-specific and developmental stage expression data where available, with direct links to source experiments. The integration covers nearly 300,000 sequences in RNAcentral. This visualisation is developed and maintained by the Expression Atlas team at EMBL-EBI. We reuse it to allow users easy access to browse and visualize the expression of RNAs in a well tested and familiar tool as shown in Figure 6A.

### API and Programmatic Access

RNAcentral continues to provide programmatic access to our data and services. As previously, we have an FTP site with data dumps, API for programmatic access, and publicly accessible postgres database. Since our last publication we have added a Swagger UI to our API documentation at: https://rnacentral.org/api/schema/swagger-ui/. We note that LitSumm summaries are also now available via the API. We now provide complete PostgreSQL database dumps via pg_dump, updated each release and compressed to approximately 42 Gb. These dumps enable local installations and high-throughput analyses without API rate limits. We would like to emphasise that in the future it may be necessary to remove old dumps if the overall size grows too large, we will only commit to providing the previous releases data dump. We encourage any users who are interested in our data at scale, particularly those who scrape our pages to use this resource.

Additionally, our services provide APIs for programmatic use. For example, sequence search and R2DT both allow automated submissions. Interested parties may find the documentation in our help pages.

Finally, RNAcentral provides a series of embeddable widgets for our services, including the sequence search, R2DT, and LitScan. These widgets have been embedded into several external resources including Rfam (LitScan, sequence search), FlyBase (R2DT), GtRNAdb (sequence search), NAKB (R2DT), PomBase (R2DT) Ribocentre, Ribocentre-switch (Sequence search), SGD (R2DT). Documentation for how to integrate resources is available in our help pages and we invite any interested resources to contact us (https://rnacentral.org/contact) to us to discuss integrating our widgets in their site.

### Licensing and Data Accessibility

In release 20, RNAcentral transitioned to a CC0 license. The CC0 license allows reuse without any attribution and is considered best practice for biological knowledgebases. We welcome anyone to reuse our data as they see fit as we place our data into the public domain as much as possible. While CC0 licensing requires no attribution, we encourage citations when RNAcentral data contributes to published research, as these metrics support continued funding and development.

### Future Directions

RNAcentral plans to continue its growth in sequences and number of data types. Our focus in the near future is to integrate more useful data types, provide more tools for community use and continue to develop our gene model and enhance our gene level annotations. To support this we will expand the quality assurance pipeline, applying additional testing for coding potential using tools such as Pfam (11) at each release. Data we are particularly interested in is RNA modifications, and so we plan to integrate thousands of nCRNA transcripts with information about posttranscriptionally modified residues from MODOMICS (32), following the integration of high-throughput modification mapping data from the

Sci-ModoM database (33). One such tool will be an MCP server for LLM access to RNAcentral data. Additionally, we plan to extend LitSumm summaries to all sequences with publications. Users interested in particular data types or researchers who are interested in collaborating are encouraged to contact us at, https://rnacentral.org/contact, and discuss their needs.

## Data availability

All data are freely available at https://rnacentral.org. The data can be accessed in the FTP archive, as well as through an API and a public Postgres database (see https://rnacentral.org/help for instructions) under the CC0 license. The code is available at https://github.com/rnacentral under the Apache 2.0 license.

## Author contributions

All RNAcentral consortium members represented here contributed to the development of their respective databases and have provided new data and features to RNAcentral since last publication.

